# Modeling cetacean eDNA distribution along the Washington coast using metabarcoding from opportunistic samples and generalized additive models

**DOI:** 10.1101/2025.11.21.689289

**Authors:** Tania Valdivia-Carrillo, Megan Shaffer, Kim Parsons, Ally Im, Andrew Shelton, Eiren K. Jacobson, Abigail Wells, Ana Ramón-Laca, Krista M. Nichols, Ryan P. Kelly, Amy Van Cise

## Abstract

Effective cetacean conservation depends on robust monitoring, yet traditional visual and passive-acoustic surveys have constraints. We evaluated environmental DNA (eDNA) metabarcoding coupled with species distribution modeling (SDM) as a tool to study habitat use of cetaceans along the Washington State coast, USA. Seawater was collected at the surface and at a 50 m depth from 43 sites (86 samples) during the 2019 U.S.–Canada Integrated Ecosystem & Acoustic-Trawl Survey. A partial section of the mitochondrial control region was amplified with cetacean - specific primers, sequenced on an Illumina MiSeq, and taxonomically assigned with a curated reference database. Nine species were detected; we modelled the three most frequent: Pacific white-sided dolphin (*Lagenorhynchus obliquidens*) (10 detections), humpback whale (*Megaptera novaeangliae*) (8 detections), and Risso’s dolphin (*Grampus griseus*) (6 detections). Binomial generalized additive models related presence-absence to bathymetry, distance to shore, longitude, slope, and sea-surface temperature; model performance was assessed with stratified five-fold cross-validation. SDMs explained 17–51% of null deviance and presented high specificity (≥ 0.80). The Pacific white-sided dolphin showed the highest eDNA presence probabilities offshore, beyond the shelf break. Humpback whale eDNA presence probabilities showed hotspots along the shelf break with secondary high-probability patches in near-shore waters. Risso’s dolphin eDNA presence probabilities were elevated in offshore zones characterized by steep bathymetric gradients, particularly northwest of the sampled transect. These spatial patterns are consistent with historical visual–acoustic records, suggesting that eDNA-informed SDMs can capture cetacean habitat use. This proof of concept indicates that combining eDNA detections with flexible SDMs could provide a cost-effective, non-invasive complement to conventional surveys and may offer a scalable pathway for marine-mammal monitoring and spatial planning.

## Introduction

Cetaceans play a critical role in marine ecosystems, yet their wide-ranging and often elusive behavior makes detecting and monitoring them difficult (Kaschner et al., 2012). Understanding and predicting their distribution is crucial for effective conservation and management in the face of increasing anthropogenic pressures and environmental changes (Tobeña et al., 2016), particularly for endangered or otherwise vulnerable species (Abrahms et al., 2019; Becker et al., 2014; Breen et al., 2017; Derville et al., 2018; Salvadeo et al., 2010; Szesciorka et al., 2023).

Species distribution models (SDMs) have become valuable tools for linking species occurrences with environmental variables, providing empirical descriptions and spatial predictions of species-environment relationships (Elith & Leathwick, 2009). For cetaceans along the US West Coast, SDMs based on systematic visual and acoustic surveys since 1991, have yielded estimates of detection, abundance, or presence probabilities by incorporating relevant environmental variables obtained from ocean models (Becker, Carretta, et al., 2020; Becker et al., 2010, 2014, 2016, 2017; Becker, Forney, et al., 2020; Fiedler et al., 2023). Within this region, the Washington State coast, part of the California Current Ecosystem (CCE), hosts a diverse array of cetaceans, including several species protected under the Endangered Species Act (United States, 1983). The CCE is characterized by a broad continental shelf, shallow basins and a strong eastern-boundary current that together create high productivity near the shelf and slope (Phillips et al., 2023), underpinning a rich marine food web that supports these cetaceans (Rice et al., 2021). Commercial shipping, fishing and naval exercises, however, pose risks to their populations (Evans, 1996; Izquierdo-Serrano et al., 2022; Lewin et al., 2019; Pirotta et al., 2019; Tixier et al., 2021). Although marine protected areas and speed-reduction zones have been implemented to mitigate such impacts, evaluating their effectiveness for these highly mobile species requires a thorough understanding of cetacean distribution patterns and habitat use (Bearzi, 2012; Nisi et al., 2024; Sahri et al., 2021).

Monitoring these populations has relied primarily on visual and acoustic surveys, yet both approaches face spatial and temporal constraints. Visual surveys provide fine spatial resolution but depend on favorable weather conditions, specialized vessels, trained observers, and they detect animals only when they surface within a limited viewing radius (Pikesley et al., 2012; Vukelic et al., 2018). Passive acoustic methods can provide continuous temporal coverage but are limited to vocally active species with described vocal repertoires (Kowarski et al., 2021) and are often affected by ambient noise (Griffiths & Barlow, 2016). These limitations underscore the need for alternative or complementary methods for cetacean monitoring.

Environmental DNA (eDNA) metabarcoding offers a powerful, non-invasive approach for cetacean monitoring by detecting species-specific genetic signatures in seawater. After quality-filtering, sequence reads are classified into amplicon sequence variants (ASVs), which capture single-nucleotide differences comparable to haplotypes (Callahan et al., 2017; Porter & Hajibabaei, 2020). ASVs are then matched against reference databases to assign taxonomy, a workflow that has proved effective for a wide range of marine taxa (Poyntz-Wright et al., 2024; Stat et al., 2018), including cetaceans (Boldrocchi et al., 2024; Merten et al., 2023; Visser et al., 2021). Cetacean eDNA detections from depths exceeding 1000 m further demonstrate eDNA’s ability to study their habitat use throughout the water column. When coupled with SDMs, these occurrence data link eDNA detections to environmental gradients, enabling prediction of habitat use over unsampled areas.

Here, we combine targeted eDNA metabarcoding with generalized additive models (GAMs)-based SDMs to map and predict the spatial distribution of cetacean eDNA along the Washington State coast. Using seawater samples collected during a large-scale systematic fisheries survey, we i) identify cetacean species present, ii) relate eDNA detection probabilities to environmental covariates via GAM-based SDMs, and iii) produce spatially explicit predictions for the most frequently detected taxa.

## Material and Methods

### eDNA Sampling

Water collection followed the fixed-station plan of the 2019 U.S.-Canada Integrated Ecosystem & Acoustic-Trawl Survey for Pacific hake (de Blois, 2020). The 43 stations span 46.40° - 48.90° N and −124.20° - −125.57° W, covering a coast-to-slope bathymetric gradient (inshore, shelf, slope) and the latitudinal range of Wahington’s continental shelf. Although opportunistic, this sampling scheme provided systematic spatial coverage of the regions’ cetacean habitats.

At each station we collected 2 samples, surface (0 m) and 50 m, for a total of 86 samples. Sampling was carried out aboard the NOAA Ship *Bell M. Shimada* between 6 and 19 August, at night to coincide with existing sampling protocols. For surface samples 2.5 L of seawater were collected from the ship’s saltwater intake; 50 m samples were taken with a Niskin bottle on a conductivity, temperature, and depth (CTD) rosette. Samples were filtered immediately through sterile mixed cellulose ester filter membranes (1 μm pore size, 47 mm-diameter) using a vacuum pump. Filters were stored in Longmire’s buffer at room temperature and later DNA was extracted following a modified phenol: chloroform method to increase yield (Ramón-Laca et al., 2021; Sambrook et al., 1989). Detailed protocols for seawater collection, preservation, and eDNA extraction are provided in Ramón-Laca et al. (2021).

### Library preparation

We performed library preparation and sequencing in triplicate for each sample. We used a 2-step PCR protocol with Dlp1.5-H and Oordlp4 primers (Baker et al., 2018) to target a 300-360 bp segment of the cetacean mitochondrial control region. The first PCR amplification was conducted using these primers with Illumina adapter overhang sequences, following the NEB Phusion HiFi Taq protocol and cycling conditions (New England Biolabs). Primers and adapter sequences are detailed in the Supplementary Material (**Table S1)**. Each 18 μl PCR1 reaction included 9 μL 2X Phusion Master Mix, 0.9 μL of each primer, 0.54 μL DMSO, 0.45 μL BSA, 4.44 μL nuclease-free water, and 1.77 μL of eDNA. The following cycling conditions were initial denaturation of 98°C for 30 sec, followed by 45 cycles of denaturation (98°C for 10 sec), annealing (60°C for 30 sec), and extension (72°C for 45 sec), with a final extension of 72°C for 10 min. After PCR amplification, products were cleaned with 1X Agencourt Ampure XP Beads (Beckman Coulter), visualized on agarose gels, and quantified using DS High Sensitivity Assay Kit (Qubit). Based on DNA quantity, an equal sample mass of the PCR product was added to the indexing PCR reaction.

The second PCR round was performed to index samples uniquely, attaching a ten-base pairs tag with a P5 or P7 MiSeq flow cell adapter via the Nextera overhang (**Table S1**), using IDT for Illumina DNA/RNA UD indexes. Each 25 μl PCR2 reaction had 12.5 μL KAPA Hifi HotStart ReadyMix, 1.25 μL of the unique IDT for Illumina DNA/RNA UD Index (one per sample), and up to 11.25 μL of the bead-cleaned PCR1 product (volume adjusted based on normalization, with the remaining volume filled with nuclease-free water). The cycling conditions were initial denaturation of 95°C for 5 min, followed by eight cycles of denaturation (98°C for 20 sec), annealing (56°C for 30 sec), and extension (72°C for 1 min), with a final extension of 72°C for 5 min. Following the index reaction, products were cleaned with 0.8X Agencourt Ampure XP beads. The final bead-cleaned product was quantified using the DS High Sensitivity Assay Kit (Qubit) and normalized to ensure equimolarity in the final library pool.

Libraries were sequenced on the Illumina MiSeq (v3 kit, 2×300 bp) at 7 pM with a 20% PhiX spike-in at the University of Washington. Each of the three technical replicates was run separately. A negative template control was included in each PCR1 to monitor contamination; all controls were sequenced alongside field samples.

### Contamination control

To minimize cross-contamination, we spatially separated pre-PCR, extraction, and post-PCR workflows in different laboratories. Workbenches were decontaminated before and after each session with DNA AWAY or 10% bleach solution followed by 70% ethanol. All pipettes and consumables (plates, tips, and tubes) were exposed to UV light for 15 minutes before use. A positive control template was processed in parallel to monitor for potential inhibition.

### Bioinformatic Analysis

Raw sequence processing was performed in R and RStudio (R Core Team, 2021; RStudio Team, 2020) using custom pipeline. Primer removal was carried out using cutadapt v1.18 (Martin, 2011). Pair-end reads were trimmed, filtered, and merged using DADA2 (Callahan et al., 2016), with truncation lengths chosen by a sliding window quality (Phred ≥29 forward, ≥30 reverse) and a minimum overlap of 42 bp for merging paired-end sequences.

Taxonomy was assigned to D-loop ASVs using DADA2’s naïve Bayesian classifier (*assignTaxonomy*) with a 98% bootstrap threshold (Wang et al., 2007), against a custom reference database including sequences from Baker et al. (2003).

After ASV taxonomic assignment, we collapsed counts to species-level read tallies. We then binarized detections by scoring a site as “present” when at least 1 read remained after all quality filters. We chose this ≥ 1-read rule to maximize sensitivity to rare taxa while stringent negative controls, chimera removal and high Phred scores minimizing false positives. The positive detections from three technical replicates and 2 depths were aggregated into a table for each species at each sampling location. We focused subsequent SDM analyses on the three most frequently detected cetacean species: Pacific white-sided dolphin, humpback whale, and Risso’s dolphin.

### Environmental predictors

Based on prior research in CCE (Becker, Carretta, et al., 2020; Becker et al., 2017; Fiedler et al., 2023), we selected five predictors: latitude, longitude, bathymetric data, distance to shore, slope, and the average seawater temperature (SST) for August 2019. Bathymetry, distance-to-shore, slope, latitude, and longitude rasters were sourced from MARSPEC (Sbrocco & Barber, 2013) via the *sdmpredictors* R package in RStudio (Bosch & Fernandez, 2023; R Core Team, 2021; RStudio Team, 2020). SST was obtained from Copernicus Global Ocean Physics Reanalysis (2023). All raster layers were standardized to 0.083° grid and clipped to the Washington State coastal polygon in QGIS (Qgis.org, 2024). Detailed variable descriptions can be found in Supplementary Material **S2**.

We then built a raster brick employing the *raster* package in R and RStudio (Hijmans, 2024; R Core Team, 2021; RStudio Team, 2020) and extracted values for each sampling site coordinate. These covariate values were subsequently used to model the probability of eDNA occurrence of individual cetacean species.

### Cetacean species distribution models (SDMs) and cross-validation from eDNA detection data

We modeled the probability of cetacean eDNA presence using binomial generalized additive models (GAMs), which flexibly relate predictor variables to binary response data via smooth functions (Wood, 2017). We fitted each smooth with thin plate regression splines (Wood, 2003) to capture nonlinear effects of bathymetry, distance to shore, slope, and seawater temperature (SST). The model for site *j* is:

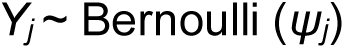

In the model, *Y_j_* is a binary indicator of eDNA presence (1) or absence (0) at site *j*, and *ψ_j_* is the probability of detection. We linked *ψ_j_* to environmental covariates using a logistic function:

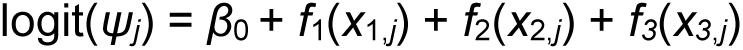

where *β*_0_ is the intercept, and each *f*_i_ is a smooth thin-plate regression spline capturing potential nonlinear effects of predictor *x_i_*at site *j*.

### Model selection and goodness-of-fit assessment

We fitted GAMs in R and RStudio (Kuhn, 2008; R Core Team, 2021; RStudio Team, 2020) using the *mgcv* and *MuMIn* packages. Each model incorporated 2-3 predictors, ensuring distance to shore and bathymetry did not co-occur (r> 0.8). We compared models using Akaike’s Information Criteria (AIC) and Bozdogan’s index of informational complexity (ICOMP).

Model fit was evaluated via adjusted R^2^ and percent deviance explained. We used *gam.check*() function in the *mgcv* package (Wood, 2023) to inspect residuals and to verify that the basis dimension for each spline was adequate using the k-index diagnostic and residual variance tests (Wood, 2017 section 5.9). All GAMs were fitted with restricted maximum likelihood (REML) and the outer Newton optimizer, converging when changes in the REML criterion fell below a predetermined threshold. Final predictions for each species were generated from the best-supported model. The code for our modeling framework can be found on GitHub.

### Cross-validation and performance evaluation

We assessed model performance with stratified k-fold cross-validation approach using the *caret* package in R and RStudio (Kuhn, 2008; R Core Team, 2021; RStudio Team, 2020). We divided the dataset into five folds (k = 5), each preserving the overall proportion of presences and absences to ensure performance estimates for the dataset. In each iteration, one-fold served as the test set and the remaining four as the training dataset. This approach was chosen because stratification better preserves the proportion of rare presences in each fold, providing more stable estimates of model performance in small, imbalanced datasets.

After obtaining probability predictions with the GAMs models for each target species, we converted them to binary presence/absence using a cut-off value of 0.3. We explored thresholds from 0.1 to 0.5 and selected 0.3 because it maximized sensitivity while keeping false-positive rate low, a priority given the small sample size and the low prevalence of detections. For each fold of the stratified five-fold cross-validation, we calculated threshold-dependent metrics (i.e., accuracy, sensitivity, specificity, and Cohen’s κ) were computed via caret’s confusion matrix and the threshold-independent area under the receiver operating characteristic curve (AUC) with the *ROCR* package in R and RStudio (R Core Team, 2021; RStudio Team, 2020; Sing et al., 2005). We report the mean ± SD of each metric across the five folds as our final performance estimates.

## Results

### Sequencing results

We successfully amplified and sequenced 86 eDNA samples at 43 stations (0 m and 50 m), conducting three independent MiSeq runs. We generated an average of 9,286,025 reads across three replicate MiSeq runs (range: 8,534,845 - 9,857,378). Table **S3** in the Supplementary Material provides detailed read counts at each step.

We identified nine cetacean species in total (<5% of the total detected ASVs; **Table 1**), with species richness per site ranging from 0 to four species (**Fig.1**). Pacific white-sided dolphin, humpback whale, and Risso’s dolphin were the most frequently detected species (**Table 1**). In total, 24 out of 43 sites had no detections for any of the three focal species.

**Figure 1.**
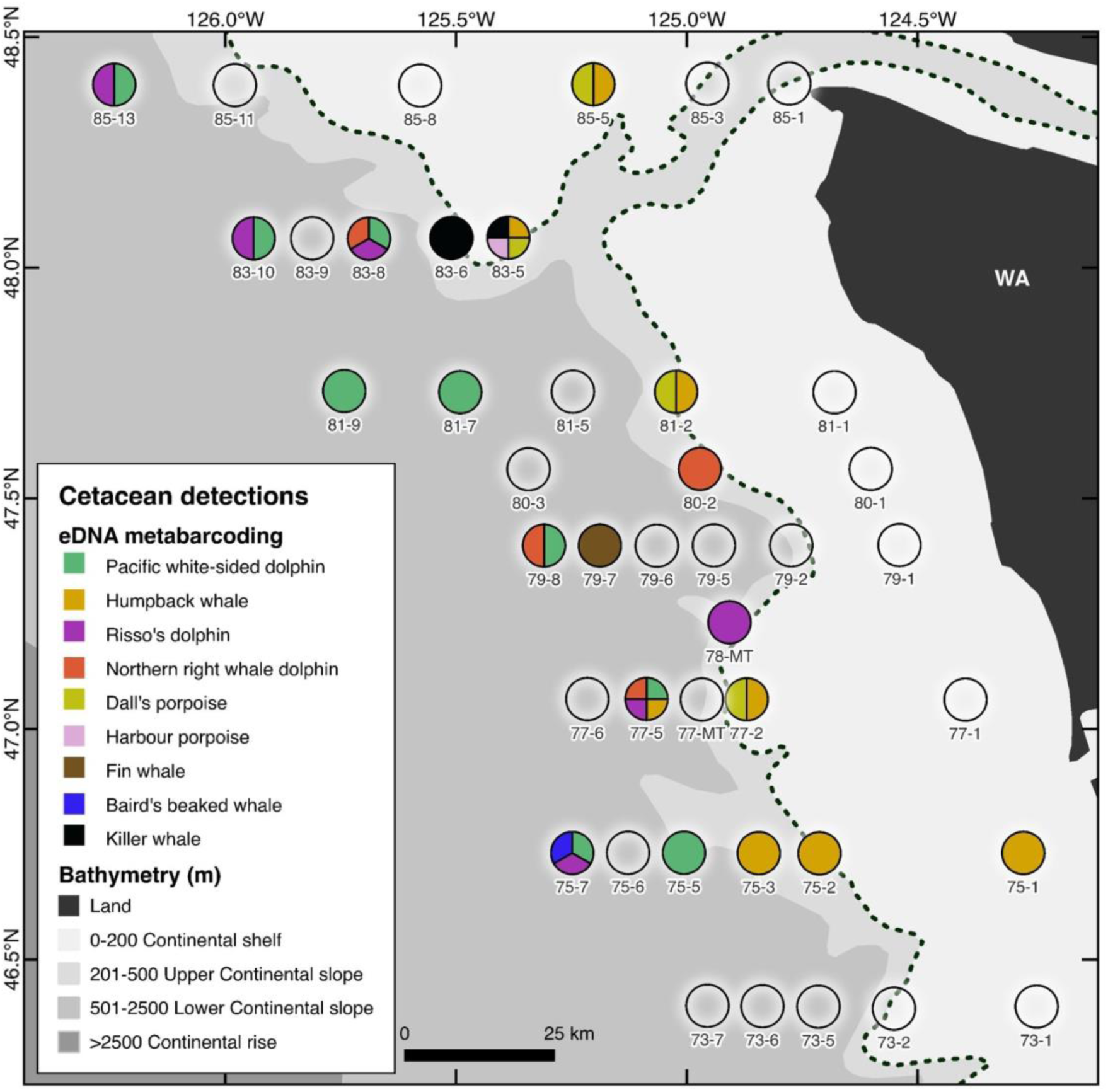
Positive cetacean detections obtained through eDNA metabarcoding of seawater samples. Open circles represent the sampling locations with no positive detections. The fill of each circle represents the species detected at that sampling location; note that data are represented as positive or negative detections for each species at each sampling location, i.e., the proportion of the circle filled by each color does not represent the relative sequence read count from that species in the sample. The dashed line is the 200 m isobath and represents the shelf break.

**Table 1.**
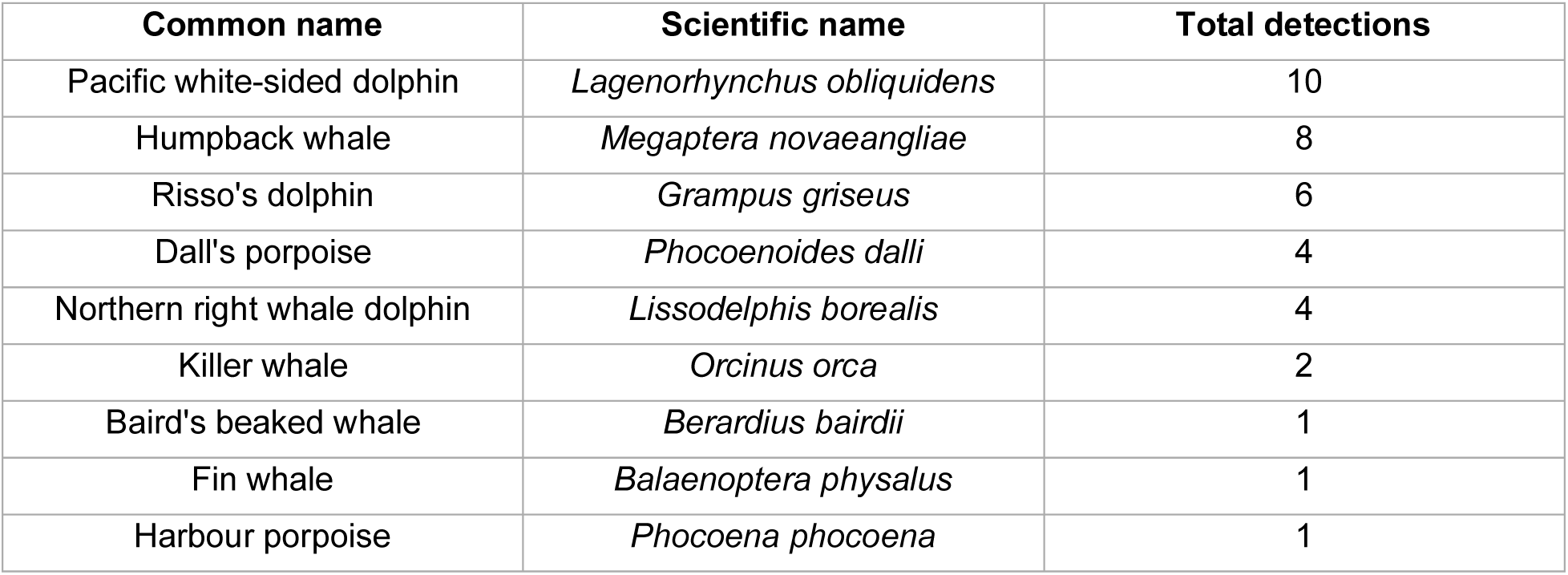
Summary of positive cetacean eDNA detections along the Washington State coast. Counts refer to independent site-depth combinations (43 sites x 2 depths). Only taxa detected at least once are shown.

| Common name | Scientific name | Total detections |
| --- | --- | --- |
| Pacific white-sided dolphin | *Lagenorhynchus obliquidens* | 10 |
| Humpback whale | *Megaptera novaeangliae* | 8 |
| Risso's dolphin | *Grampus griseus* | 6 |
| Dall's porpoise | *Phocoenoides dalli* | 4 |
| Northern right whale dolphin | *Lissodelphis borealis* | 4 |
| Killer whale | *Orcinus orca* | 2 |
| Baird's beaked whale | *Berardius bairdii* | 1 |
| Fin whale | *Balaenoptera physalus* | 1 |
| Harbour porpoise | *Phocoena phocoena* | 1 |

### Cetacean species distribution models (SDMs) from eDNA detection data

#### Pacific white-sided dolphin

The best-fitting GAM model retained two smooth predictors, longitude (*p* = 0.046) and distance to shore (*p* = 0.009), both significant (**Table 2**). Together they explained 47.7 % of the variation in eDNA presence on the link scale (adjusted R²) and reduced the null deviance by 50.6 %, indicating that half of the structure in the occurrence data is captured by these spatial variables. The model converged after five iterations, and k-index diagnostics confirmed adequate basis dimensions (k-index p_lon_ = 0.97; p_dist_shore_ = 0.95), providing no evidence of over- or under-smoothing.

**Table 2.** Best GAM models for the three more frequent cetacean species detected along the Washington coast via eDNA metabarcoding. Models incorporate different environmental variables to predict eDNA-spatial distribution, including lat (latitude), lon (longitude), bathy (bathymetry), dist_shore (distance to shore), slope, and seawater temperature (SST). Statistical output parameters include R^2^ (adjusted), and deviance explained. * p-value < 0.05

| Species | (Int) | lat | lon | bathy | dist_shore | slope | SST | R<sup>2</sup>(adj) | dev. explained |
| --- | --- | --- | --- | --- | --- | --- | --- | --- | --- |
| Pacific white-sided dolphin | 0.003* |  | 0.046* |  | 0.009* |  |  | 0.477 | 50.60 |
| Humpback whale | 0.000* |  |  | 0.003* |  |  | 0.026* | 0.285 | 29.20 |
| Risso's dolphin | 0.000* |  | 0.038* |  |  | 0.053 |  | 0.165 | 22.90 |

#### Humpback whale

In the best-fitting GAM model for the humpback whale, eDNA bathymetry (*p* = 0.003) and sea-surface temperature (*p* = 0.026) were retained as significant smooth terms (**Table 2**). These predictors explained 28.5% of the variation in eDNA presence (adjusted R²) and reduced the null deviance by 29.2%, indicating that one-third of the spatial-environmental signal is captured. Convergence was reached after four iterations, and k-index diagnostics confirmed the adequate basis dimensions (k-index p_bathy_ = 0.52 and p_SST_ = 0.095) with no evidence of over- or under-smoothing.

#### Risso’s dolphin

The best-fitting GAM model for Risso’s dolphin retained longitude (*p* = 0.038) and slope (*p* = 0.053) as smooth terms (**Table 2**). These predictors accounted for 16.5% of the variation in eDNA presence (adjusted R²) and reduced the null deviance by 22.9%, suggesting a modest yet measurable improvement over the intercept-only model (null). The fit converged stably, and k-index diagnostics confirmed adequate basis dimensions (k-index p_lon_ = 0.97; p_slope_ = 0.52), suggesting no evidence of over- or under-smoothing.

#### Spatial patterns

Projecting these models onto the study area (**Fig.2**) revealed distinct habitat associations. Pacific white-sided dolphin eDNA shows a higher probability of occurrence in offshore regions, beyond the shelf break. Humpback whale eDNA showed higher probabilities along the shelf break with localized coastal hotspots. Risso’s dolphin eDNA was most probable in offshore regions with steep bathymetric features and hotspots northwest of the sampled area.

**Figure 2.**
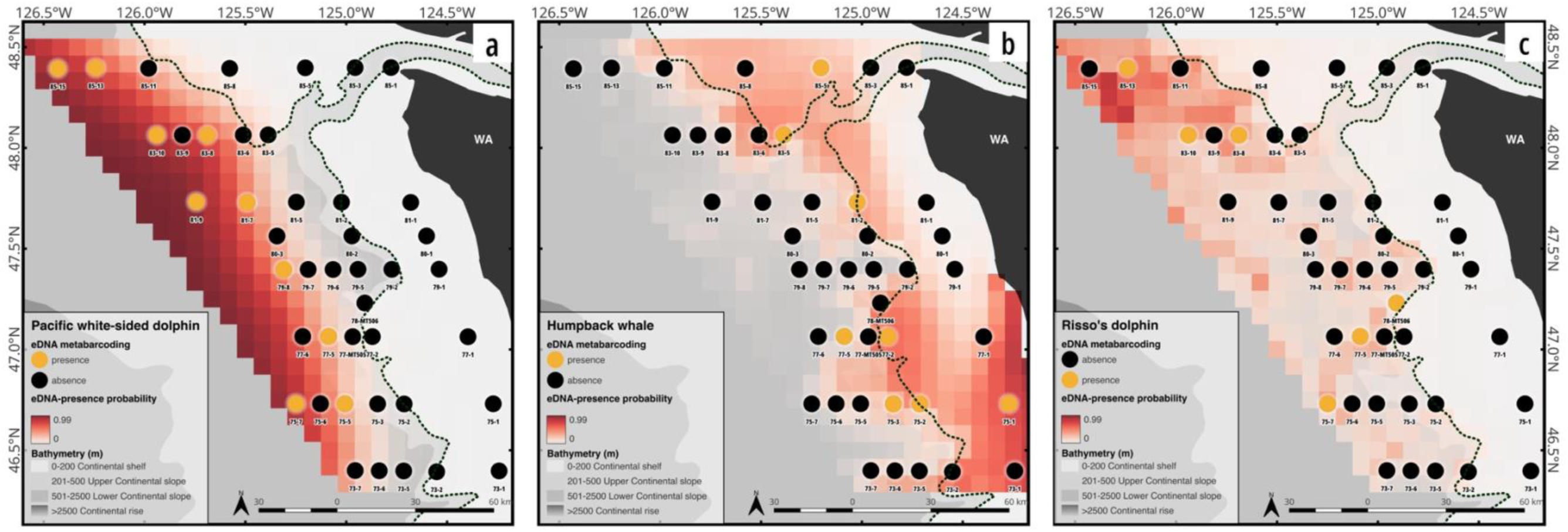
eDNA presence probability derived from GAM models to evaluate eDNA species distribution for the a) Pacific white-sided dolphin, b) Humpback whale, and c) Risso’s dolphin. The dashed line is the 200 m isobath and represents the shelf break.

**Table 2** and **Fig.2** summarize the best GAM outputs and predicted eDNA presence probabilities, respectively, illustrating an offshore distribution for Pacific white-sided dolphin and Risso’s dolphins, and shelf-break/coastal hotspots for humpback whales.

The Table **S4** and Figure **S5** of the Supplementary Material presents detailed information on the modeling results.

### Cross-validation for eDNA-based models

#### Pacific white-sided dolphin

The GAM for *Lagenorhynchus obliquidens* exhibited the highest predictive accuracy, with a mean accuracy of 0.84 ± 0.19 and an area under the ROC curve (AUC) of 0.92 ± 0.17. Cohen’s kappa (κ = 0.61 ± 0.39) indicated substantial agreement beyond chance. Sensitivity, the true-positive rate, averaged 0.75 ± 0.27, reflecting the model’s strong ability to detect actual eDNA presences. Specificity remained high at 0.87 ± 0.19, demonstrating reliable identification of absences. The F₁-score of 0.72 ± 0.27 confirms a balanced trade-off between precision and recall.

#### Humpback whale

For *Megaptera novaeangliae*, the model yielded a mean accuracy of 0.80 ± 0.17 and AUC of 0.87 ± 0.22, indicative of good overall discrimination. Cohen’s κ = 0.45 ± 0.45 denoted moderate agreement beyond chance, with sensitivity of 0.70 ± 0.45 and specificity of 0.85 ± 0.18. The F₁-score of 0.68 ± 0.25 illustrates that the model maintains a reasonable balance between detecting true positives and minimizing false positives, despite increased variability associated with fewer positive cases.

#### Risso’s dolphin

The GAM for Grampus griseus faced limitations due to a small number of positive detections (n = 6). It achieved a mean accuracy of 0.74 ± 0.15 but an AUC of only 0.62 ± 0.16, indicating modest discriminative performance. Cohen’s κ = 0.06 ± 0.32 suggests near-random agreement beyond chance. Sensitivity was low (0.30 ± 0.45), while specificity remained comparatively robust at 0.80 ± 0.16. The F₁-score of 0.50 ± 0.24 further highlights the challenge of accurately predicting rare eDNA occurrences. **Table 3** summarizes the mean ± SD of classification metrics across stratified five-fold cross-validation for each species.

**Table 3.** Summary of cross-validation results for three more frequent cetacean species detected along the Washington coast via eDNA metabarcoding (threshold = 0.3). Values are reported as mean ± SD across folds.

| Species | Accuracy | Cohen's K | Sensitivity | Specificity | F1 | AUC |
| --- | --- | --- | --- | --- | --- | --- |
| Pacific white-sided dolphin | 0.84 ± 0.19 | 0.61 ± 0.39 | 0.75 ± 0.27 | 0.87 ± 0.19 | 0.72 ± 0.27 | 0.92 ± 0.17 |
| Humpback whale | 0.8 ± 0.17 | 0.45 ± 0.45 | 0.7 ± 0.45 | 0.85 ± 0.18 | 0.68 ± 0.25 | 0.87 ± 0.22 |
| Risso's dolphin | 0.74 ± 0.15 | 0.06 ± 0.32 | 0.3 ± 0.45 | 0.8 ± 0.16 | 0.5 ± 0.24 | 0.62 ± 0.16 |

## Discussion

This study demonstrates the feasibility of integrating environmental DNA (eDNA) metabarcoding detection data into species distribution models (SDMs) to investigate cetacean spatial distributions along the Washington State coast. By applying generalized additive models (GAMs), we explored the relationship between eDNA presence and environmental variables for the three most frequently detected species: Pacific white-sided dolphin, humpback whale, and Risso’s dolphin. Model performance was confirmed with stratified five-fold cross-validation, which showed consistently high values of accuracy, AUC, κ, and other diagnostics, indicating that the eDNA-based SDMs generalize well to unseen data for the Pacific white-sided dolphin and humpback whale, whereas performance for Risso’s dolphin was limited by sparse detections.

It is important to clarify that the models predict the spatial distribution of eDNA detections, not the locations of individual cetaceans. Detected DNA could originate from animals that were present locally or from genetic material transported by currents, and the time elapsed since shedding is uncertain. Nevertheless, by mapping where cetacean genetic material is most likely to occur under prevailing environmental conditions, our approach offers a first step toward incorporating eDNA evidence into predictive frameworks for cetaceans, providing insights that can enhance conservation and management strategies.

### Cetacean SDMs from eDNA detection data

#### Pacific white-sided dolphin

Pacific white-sided dolphins are pelagic species commonly found in Washington waters during the summer months. They take advantage of the productive high-latitude waters for foraging (Forney & Barlow, 1998; Rice et al., 2021). The CCE plays a vital role in shaping their distribution, as nutrient upwelling enhances prey availability in offshore waters, particularly at the shelf break and upper slope regions (Becker et al., 2022; Rice et al., 2021). Our model confirms these findings, showing that the probability of detecting environmental DNA (eDNA) increases beyond the 200-meter isobath. This aligns with previous studies identifying the continental shelf break as a crucial habitat for dolphins (Giménez et al., 2021; Gómez-Campos et al., 2011).

#### Humpback whale

Humpback whales undertake long-distance migrations between tropical breeding grounds and high-latitude summer feeding areas, including the coast of Washington (Baker et al., 1994; Barlow et al., 2011; Darling et al., 2022; Wright et al., 2015). Their distribution is influenced by bathymetric features and highly productive upwelling zones that concentrate prey (Derville et al., 2018; Fleming et al., 2016; Garrigue et al., 2015; Thompson et al., 2012). Our model shows a strong correlation between humpback whale eDNA presence and both bathymetry and SST. This pattern is consistent with earlier studies that link bathymetry and SST to humpback foraging along shelf-break regions (Becker, Carretta, et al., 2020; Burrows et al., 2016; Dalla Rosa et al., 2012; Meynecke et al., 2021; Miller et al., 2022; Pelayo-González et al., 2022; Rockwood et al., 2020), and our predicted eDNA hotspots coincide with shelf-break areas of high krill biomass (Phillips et al., 2023), underscoring prey availability as driver of humpback whale distribution (Rockwood et al., 2020). Consistently, both acoustic (Rice et al., 2021) and visual (Barlow & Forney, 2007; Schorr et al., 2007) surveys report the highest humpback occurrence over the continental shelf along the Washington coast.

#### Risso’s dolphin

Risso’s dolphins are typically found as offshore cetaceans, often observed near the outer edges of continental shelves and upper slopes (Azzellino et al., 2008; Baumgartner, 1997; Cañadas et al., 2002; Praca & Gannier, 2008). This habitat corresponds to areas where their primary prey, cephalopods, are most abundant (Praca & Gannier, 2008; Visser et al., 2021). Our model identified a significant association between the probability of detecting Risso’s dolphin eDNA and longitude, consistent with prior research highlighting the species’ preference for offshore environments (Becker, Carretta, et al., 2020; Becker, Forney, et al., 2020). Although slope was non-significant in our model (p = 0.053), its exact role remains unclear. Higher-resolution bathymetric data could clarify whether slope influences habitat selection by affecting prey aggregation. Our results suggest that this study area may represent a transitional zone, where Risso’s dolphin presence declines at higher latitudes as water temperatures become more polar (Frouin-Mouy et al., 2022; Jefferson et al., 2014).

Both Pacific white-sided dolphins and Risso’s dolphins primarily forage on squid and exhibit nocturnal activity. Our eDNA sampling at night likely coincided with an increase in surface presence, which may enhance detection rates. This diel pattern aligns with previous studies showing that delphinids engage in vertical migration in search of their prey (Benoit-Bird et al., 2004; Benoit-Bird & Au, 2003).

#### Model robustness and limitations by cross-validation

Cross-validation serves as a critical tool for evaluating the robustness of models, particularly in scenarios involving small datasets. The outcomes of stratified k-fold cross-validation for the three focal species provide valuable insights into the predictive capacity and inherent limitations of our eDNA-based SDMs. Stratified 5-fold cross-validation showed that our eDNA-based GAMs capture meaningful spatial signal but also revealed limits imposed by a presence-rare data set. Accuracy remained ≥ 0.74 and specificity ≥ 0.80 for all three taxa, indicating few false positives. Metrics sensitive to the number of detections diverged: the Pacific white-sided dolphin model, supported by the largest set of detections, showed the strongest and most reliable discrimination (mean AUC = 0.92, κ = 0.61), whereas the humpback whale model was moderately robust (AUC = 0.87) but variable across folds, reflecting data scarcity. With only six positive sites, the Risso’s dolphin model kept high specificity yet low sensitivity (mean 0.30) and near-random agreement (κ = 0.06), highlighting the difficulty of modeling sparse eDNA records. Taken together, these results suggest that eDNA-SDMs built from ≥ 10 detections provide reproducible spatial inference; below that threshold, uncertainty increases. Also, even when presence data are limited, cross-validation diagnostics, particularly sensitivity and κ, offer an objective benchmark for deciding whether predictions are fit for management use or should be treated as exploratory.

#### Sampling-design and threshold considerations

All eDNA samples were collected opportunistically at the fixed stations of the Integrated Ecosystem & Acoustic-Trawl Survey, which ensured a broad latitudinal and bathymetric coverage but did under-represented shallow embayment (< 50 m) and outer-shelf waters. This sampling may bias SDM predictions toward the environmental envelope of the survey transects. Additionally, we adopted sensitive detection rules, both in the laboratory (≥ 1 read = presence) and in model evaluation (0.30 probability cut-off), to minimize false negatives given the low detection frequencies. Although contamination controls mitigate false positives, future studies should integrate targeted eDNA stations in unsampled coastal and offshore regions and increase biological replication to reduce spatial bias and increase detections.

#### Challenges of using eDNA data in SDMs

The interpretation of eDNA read counts presents several methodological and ecological challenges that complicate their direct use in species distribution models (SDMs). While eDNA metabarcoding enables species detection through high-throughput sequencing, read counts do not necessarily reflect organism abundance due to various biological, technical, and environmental factors. Given these complexities, we opted to transform our eDNA data into a presence-absence format to improve model robustness and interpretability.

eDNA-based methods, though highly sensitive, share limitations with traditional monitoring approaches (Fukaya et al., 2022; Mathieu et al., 2020). The amplification of specific DNA molecules can be influenced by stochastic processes, including the limited availability of starting material, which may lead to variability in detection outcomes (Kelly et al., 2019). Additionally, detection probability is significantly affected by environmental conditions, sampling design, and replication, particularly for rare or low-density species (Lahoz-Monfort et al., 2016). To minimize underrepresentation of rare species, it is advisable to increase both biological and technical replicates, as previous studies have demonstrated that replication enhances detection rates and reduces bias (Alberdi et al., 2018; Buxton et al., 2021; Ficetola et al., 2015). The lack of biological replication in this study may contribute to an underestimation of rare species’ presence.

eDNA persistence and detectability in the environment vary due to multiple biological, physical, and ecological factors (Brasseale et al., 2025). Shedding rates differ substantially across species and are influenced by physiological and behavioral traits. Cetaceans, for example, release eDNA through sloughed skin and feces, but the quantity shed is highly dependent on life-history traits, body size, and health condition. Moreover, species-specific shedding rates fluctuate with seasonality and behavior (Allan et al., 2020; Collins et al., 2018), further complicating the interpretation of eDNA concentrations as direct indicators of abundance. Once released, eDNA undergoes degradation at variable rates depending on environmental conditions such as temperature, UV exposure, microbial activity, and water chemistry (Barnes et al., 2014). In cases where degradation outpaces accumulation, the presence of a species may be underestimated, while in other cases, eDNA may persist longer than expected, leading to false positives if not carefully accounted for.

Another key consideration is the transport of eDNA in marine environments. Unlike terrestrial systems, marine eDNA can be dispersed by currents, tides, and vertical mixing (Allan et al., 2021; Andruszkiewicz et al., 2019; Brasseale et al., 2025). This movement complicates the interpretation of read counts because high eDNA concentrations at a given location may not necessarily indicate local species presence but rather the accumulation of transported genetic material from upstream sources. Given this spatial complexity, a presence-absence framework provides a more conservative and ecologically meaningful approach to SDMs.

As the scientific community improves its understanding of eDNA dynamics, we anticipate advances in species detection accuracy, habitat modeling, and SDM precision. Further developments, such as quantifying eDNA concentrations and linking them to species abundance (e.g., Guri et al., 2024), could significantly enhance model performance and ecological inference. The integration of eDNA data with comprehensive environmental datasets, including hydrodynamic models, could allow for more precise estimations of species distributions while accounting for eDNA transport and degradation processes.

Our study illustrates that eDNA metabarcoding can serve as a stand-alone tool for SDM applications, particularly in scenarios where traditional survey methods (e.g., visual and acoustic surveys) are impractical or resource intensive. While traditional observational datasets exist for cetaceans in our study area (e.g., Becker et al., 2014), our research underscores the feasibility of using eDNA metabarcoding to detect and model cetacean presence in a cost-effective, non-invasive manner. However, this study serves as a proof of concept, and its statistical power is inherently constrained by a small sample size. The continued evolution of eDNA methodologies, combined with advancements in oceanographic modeling and environmental DNA transport simulations, presents a clear opportunity for refining and expanding SDM applications. By incorporating biotic and abiotic environmental predictors, as well as improved molecular techniques, these models can be extended to a wide range of marine species, enhancing their utility for conservation and management.

## Conclusions

Our results demonstrate that environmental DNA metabarcoding, coupled with generalized additive modeling, can generate ecologically coherent species-distribution estimates for highly mobile cetaceans. The study was intentionally exploratory, with 43 stations and a limited number of positive detections per species (1–10 occurrences) inherently reduce statistical power and model generalizability. Even so, stratified k-fold cross-validation confirmed that models built from as few as 10 positive sites can deliver reproducible spatial signal, whereas models with fewer detections should be regarded as preliminary. The lower sensitivity observed for Risso’s dolphins underscores the challenges associated with modeling species with sparse detections, emphasizing the need for increased sampling efforts across broader spatial and temporal scales. A key challenge in eDNA-based SDMs is that they predict areas where cetacean DNA was detected, rather than direct species presence. This distinction is particularly important in marine systems, where eDNA transport via ocean currents, vertical mixing, and degradation can influence detection locations. Future research should integrate hydrodynamic modeling to better account for these transport processes, thereby improving spatial inferences of cetacean distribution. Despite these challenges, the integration of eDNA detection data into SDMs presents a significant opportunity for advancing cetacean monitoring and conservation. By expanding sampling efforts, incorporating hydrodynamic models, and refining molecular methodologies, future studies can enhance the accuracy and ecological relevance of eDNA-based distribution models. This research sets the stage for the broader adoption of eDNA metabarcoding in marine conservation, providing cost-effective, scalable, and non-invasive monitoring solutions for cetaceans and other marine megafauna.

## Data Archiving Statement

Raw fastq files will be uploaded to NCBI-Genbank and will be ready for publication. The code used for our analysis is available on GitHub.

## Author contribution

The collection of the samples was supported by the NOAA Fisheries Genomics Strategic Initiative, led by KN. TVC, RK, AVC, and AS developed the ideas and designed the methodology for this study. AW and ARL collected the eDNA samples and performed DNA extractions. MS conducted laboratory experiments. AVC, AI, and TVC constructed the bioinformatic pipeline. TVC analyzed the data obtained from bioinformatic analysis and led the manuscript writing. All authors critically reviewed and contributed to the drafts and provided final approval for publication.

## Data Archiving Statement

Raw fastq files will be uploaded to NCBI-Genbank and will be ready for publication. The code used for our analysis is available in the GitHub repositories: https://github.com/MMARINeDNA/metabarcoding_QAQC_pipeline and https://github.com/MMARINeDNA/Valdiviaetal_2D-eDNA-spatial-distributions/tree/main, under the CC0-1.0 license.

## Acknowledgments

The authors gratefully acknowledge the survey expertise and support of NOAA’s Pacific Hake survey team and the NOAA Ship Bell M. Shimada personnel for supporting the collection of eDNA samples on the West Coast Hake Survey 2019-2023.

## References

Abrahms, B., Welch, H., Brodie, S., Jacox, M. G., Becker, E. A., Bograd, S. J., Irvine, L. M., Palacios, D. M., Mate, B. R., & Hazen, E. L. (2019). Dynamic ensemble models to predict distributions and anthropogenic risk exposure for highly mobile species. Diversity & Distributions, 25(8), 1182–1193.

Alberdi, A., Aizpurua, O., Gilbert, M. T. P., & Bohmann, K. (2018). Scrutinizing key steps for reliable metabarcoding of environmental samples. 2018(May 2017), 134–147.

Allan, E. A., DiBenedetto, M. H., Lavery, A. C., Govindarajan, A. F., & Zhang, W. G. (2021). Modeling characterization of the vertical and temporal variability of environmental DNA in the mesopelagic ocean. Scientific Reports, 11(1), 21273.

Allan, E. A., Zhang, W., Lavery, A., & Govindarajan, A. F. (2020). Environmental DNA shedding and decay rates from diverse animal forms and thermal regimes. 10.1002/EDN3.141

Andruszkiewicz, E. A., Koseff, J. R., Fringer, O. B., Ouellette, N. T., Lowe, A. B., Edwards, C. A., & Boehm, A. B. (2019). Modeling Environmental DNA Transport in the Coastal Ocean Using Lagrangian Particle Tracking. Frontiers in Marine Science, 6. 10.3389/fmars.2019.00477

Azzellino, A., Gaspari, S., Airoldi, S., & Nani, B. (2008). Habitat use and preferences of cetaceans along the continental slope and the adjacent pelagic waters in the western Ligurian Sea. Deep Sea Research Part I: Oceanographic Research Papers, 55(3), 296–323.

Baker, C. S., Slade, R. W., Bannister, J. L., Abernethy, R. B., Weinrich, M. T., Lien, J., Urban, J., Corkeron, P., Calmabokidis, J., & Vasquez, O. (1994). Hierarchical structure of mitochondrial DNA gene flow among humpback whales Megaptera novaeangliae, world-wide. Molecular Ecology, 3(4), 313–327.

Baker, C. S., Steel, D., Nieukirk, S., & Klinck, H. (2018). Environmental DNA (eDNA) From the Wake of the Whales: Droplet Digital PCR for Detection and Species Identification. Frontiers in Marine Science, 5. 10.3389/fmars.2018.00133

Baker, S. C., Lavery, D. M. L., & Shane, R. H. A. (2003). www.DNA-surveillance: applied molecular taxonomy for species conservation and discovery. Trends in Ecology & Evolution, 18(6), 271–272.

Barlow, J., Calambokidis, J., Falcone, E. A., Baker, C. S., Burdin, A. M., Clapham, P. J., Ford, J. K. B., Gabriele, C. M., LeDuc, R., Mattila, D. K., Quinn, T. J., II, Rojas-Bracho, L., Straley, J. M., Taylor, B. L., Urbán R., J., Wade, P., Weller, D., Witteveen, B. H., & Yamaguchi, M. (2011). Humpback whale abundance in the North Pacific estimated by photographic capture-recapture with bias correction from simulation studies. Marine Mammal Science, 27(4), 793–818.

Barlow, J., & Forney, K. (2007). Abundance and population density of cetaceans in the California Current ecosystem. https://aquadocs.org/handle/1834/25509

Barnes, M. A., Turner, C. R., Jerde, C. L., Renshaw, M. A., Chadderton, W. L., & Lodge, D. M. (2014). Environmental conditions influence eDNA persistence in aquatic systems. Environmental Science & Technology, 48(3), 1819–1827.

Baumgartner, M. F. (1997). The distribution of Risso’s dolphin (*Grampus griseus*) with respect to the physiography of the northern gulf of Mexico. Marine Mammal Science, 13(4), 614–638.

Bearzi, M. (2012). Cetaceans and MPAs should go hand in hand: A case study in Santa Monica Bay, California. Ocean & Coastal Management, 60, 56–59.

Becker, E. A., Carretta, J. V., Forney, K. A., Barlow, J., Brodie, S., Hoopes, R., Jacox, M. G., Maxwell, S. M., Redfern, J. V., Sisson, N. B., Welch, H., & Hazen, E. L. (2020). Performance evaluation of cetacean species distribution models developed using generalized additive models and boosted regression trees. Ecology and Evolution, 10(12), 5759–5784.

Becker, E. A., Forney, K. A., Ferguson, M. C., Foley, D. G., Smith, R. C., Barlow, J., & Redfern, J. V. (2010). Comparing California Current cetacean–habitat models developed using in situ and remotely sensed sea surface temperature data. Marine Ecology Progress Series, 413, 163–183.

Becker, E. A., Forney, K. A., Fiedler, P. C., Barlow, J., Chivers, S. J., Edwards, C. A., Moore, A. M., & Redfern, J. V. (2016). Moving Towards Dynamic Ocean Management: How Well Do Modeled Ocean Products Predict Species Distributions? Remote Sensing, 8(2), 149.

Becker, E. A., Forney, K. A., Foley, D. G., Smith, R. C., Moore, T. J., & Barlow, J. (2014). Predicting seasonal density patterns of California cetaceans based on habitat models. Endangered Species Research, 23(1), 1–22.

Becker, E. A., Forney, K. A., Miller, D. L., Barlow, J., Rojas-Bracho, L., Urbán R, J., & Moore, J. E. (2022). Dynamic Habitat Models Reflect Interannual Movement of Cetaceans Within the California Current Ecosystem. Frontiers in Marine Science, 9. 10.3389/fmars.2022.829523

Becker, E. A., Forney, K. A., Miller, D. L., Fiedler, P. C., Barlow, J., & Moore, J. E. (2020). Habitat-based density estimates for cetaceans in the California Current Ecosystem based on 1991-2018 survey data. https://repository.library.noaa.gov/view/noaa/27826

Becker, E. A., Forney, K. A., Thayre, B. J., Debich, A. J., Campbell, G. S., Whitaker, K., Douglas, A. B., Gilles, A., Hoopes, R., & Hildebrand, J. A. (2017). Habitat-Based Density Models for three Cetacean Species off Southern California Illustrate Pronounced Seasonal Differences. Frontiers in Marine Science, 4. 10.3389/fmars.2017.00121

Benoit-Bird, K. J., & Au, W. W. L. (2003). Prey dynamics affect foraging by a pelagic predator (Stenella longirostris) over a range of spatial and temporal scales. Behavioral Ecology and Sociobiology, 53(6), 364–373.

Benoit-Bird, K. J., Würsig, B., & Mfadden, C. J. (2004). Dusky dolphin (*Lagenorhynchus obscurus*) foraging in two different habitats: active acoustic detection of dolphins and their prey. Marine Mammal Science, 20(2), 215–231.

Boldrocchi, G., Conte, L., Galli, P., Bettinetti, R., & Valsecchi, E. (2024). Cuvier’s beaked whale (Ziphius cavirostris) detection through surface-sourced eDNA: A promising approach for monitoring deep-diving cetaceans. Ecological Indicators, 161(111966), 111966.

Bosch, S., & Fernandez, S. (2023). sdmpredictors: Species Distribution Modelling Predictor Datasets (Version 0.2.15) [R package]. http://lifewatch.github.io/sdmpredictors/

Brasseale, E., Adams, N., Allan, E. A., Jacobson E. K., Kelly, R. P., Liu, O. R., Moore, S., Shaffer, M., Xiong, J. and Parsons, K. 2025. Marine eDNA Production and Loss Mechanisms. Journal of Geophysical Research. Oceans 130 (4): e2024JC021643.

Breen, P., Brown, S., Reid, D., & Rogan, E. (2017). Where is the risk? Integrating a spatial distribution model and a risk assessment to identify areas of cetacean interaction with fisheries in the northeast Atlantic. Ocean & Coastal Management, 136, 148–155.

Burrows, J. A., Johnston, D. W., Straley, J. M., Chenoweth, E. M., Ware, C., Curtice, C., DeRuiter, S. L., & Friedlaender, A. S. (2016). Prey density and depth affect the fine-scale foraging behavior of humpback whales Megaptera novaeangliae in Sitka Sound, Alaska, USA. Marine Ecology Progress Series, 561, 245–260.

Buxton, A., Matechou, E., Griffin, J., Diana, A., & Griffiths, R. A. (2021). Optimising sampling and analysis protocols in environmental DNA studies. Scientific Reports, 11(1), 11637.

Callahan, B. J., McMurdie, P. J., & Holmes, S. P. (2017). Exact sequence variants should replace operational taxonomic units in marker-gene data analysis. The ISME Journal, 11(12), 2639–2643.

Callahan, B. J., McMurdie, P. J., Rosen, M. J., Han, A. W., Johnson, A. J. A., & Holmes, S. P. (2016). DADA2: High-resolution sample inference from Illumina amplicon data. Nature Methods, 13(7), 581–583.

Cañadas, A., Sagarminaga, R., & Garcı’a-Tiscar, S. (2002). Cetacean distribution related with depth and slope in the Mediterranean waters off southern Spain. Deep Sea Research Part I: Oceanographic Research Papers, 49(11), 2053–2073.

Collins, R. A., Wangensteen, O. S., Gorman, E. J. O., Mariani, S., Sims, D. W., & Genner, M. J. (2018). Persistence of environmental DNA in marine systems. Communications Biology, December. 10.1038/s42003-018-0192-6

Dalla Rosa, L., Ford, J. K. B., & Trites, A. W. (2012). Distribution and relative abundance of humpback whales in relation to environmental variables in coastal British Columbia and adjacent waters. Continental Shelf Research, 36, 89–104.

Darling, J. D., Audley, K., Cheeseman, T., Goodwin, B., Lyman, E. G., & Urbán, R. J. (2022). Humpback whales (Megaptera novaeangliae) attend both Mexico and Hawaii breeding grounds in the same winter: mixing in the northeast Pacific. Biology Letters, 18(2), 20210547.

de Blois, S. (2020). The 2019 joint U.s.-Canada integrated ecosystem and pacific hake acoustic-trawl survey: Cruise report SH-19-06. Northwest Fisheries Science Center (U.S.). 10.25923/P55A-1A22

Derville, S., Torres, L. G., Iovan, C., & Garrigue, C. (2018). Finding the right fit: Comparative cetacean distribution models using multiple data sources and statistical approaches. Diversity and Distributions, 24(11), 1657–1673.

Elith, J., & Leathwick, J. R. (2009). Species Distribution Models: Ecological Explanation and Prediction Across Space and Time. Annual Review of Ecology, Evolution, and Systematics, 40(1), 677–697.

Evans, P. G. H. (1996). Human disturbance of cetaceans. In The Exploitation of Mammal Populations (pp. 376–394). Springer Netherlands.

Ficetola, G. F., Pansu, J., Bonin, A., Coissac, E., Giguet-Covex, C., De Barba, M., Gielly, L., Lopes, C. M., Boyer, F., Pompanon, F., Rayé, G., & Taberlet, P. (2015). Replication levels, false presences and the estimation of the presence/absence from eDNA metabarcoding data. Molecular Ecology Resources, 15(3), 543–556.

Fiedler, P. C., Becker, E. A., Forney, K. A., Barlow, J., & Moore, J. E. (2023). Species distribution modeling of deep-diving cetaceans. Marine Mammal Science. 10.1111/mms.13057

Fleming, A. H., Clark, C. T., Calambokidis, J., & Barlow, J. (2016). Humpback whale diets respond to variance in ocean climate and ecosystem conditions in the California Current. Global Change Biology, 22(3), 1214–1224.

Forney, K. A., & Barlow, J. (1998). Seasonal patterns in the abundance and distribution of California cetaceans, 1991–1992. Marine Mammal Science, 14(3), 460–489.

Frouin-Mouy, H., Mouy, X., Pilkington, J., Küsel, E., Nichol, L., Doniol-Valcroze, T., & Lee, L. (2022). Acoustic and visual cetacean surveys reveal year-round spatial and temporal distributions for multiple species in northern British Columbia, Canada. Scientific Reports, 12(1), 19272.

Fukaya, K., Kondo, N. I., Matsuzaki, S.-I. S., & Kadoya, T. (2022). Multispecies site occupancy modelling and study design for spatially replicated environmental DNA metabarcoding. Methods in Ecology and Evolution / British Ecological Society, 13(1), 183–193.

Garrigue, C., Clapham, P. J., Geyer, Y., Kennedy, A. S., & Zerbini, A. N. (2015). Satellite tracking reveals novel migratory patterns and the importance of seamounts for endangered South Pacific humpback whales. Royal Society Open Science, 2(11), 150489.

Giménez, J., Authier, M., Valeiras, J., Abad, E., Marçalo, A., Coll, M., Gauffier, P., Santos, M. B., & de Stephanis, R. (2021). Consumption rates and interaction with fisheries of Mediterranean common dolphins in the Alboran Sea. Regional Studies in Marine Science, 45(101826), 101826.

Global Ocean Physics Reanalysis. (2023). [Dataset]. In COPERNICUS. https://data.marine.copernicus.eu/product/GLOBAL_MULTIYEAR_PHY_001_030/description

Gómez-Campos, E., Borrell, A., Cardona, L., Forcada, J., & Aguilar, A. (2011). Overfishing of small pelagic fishes increases trophic overlap between immature and mature striped dolphins in the Mediterranean Sea. PloS One, 6(9), e24554.

Griffiths, E. T., & Barlow, J. (2016). Cetacean acoustic detections from free-floating vertical hydrophone arrays in the southern California Current. The Journal of the Acoustical Society of America, 140(5), EL399.

Guri, G., Shelton, A. O., Kelly R. P., Yoccoz, N., Johansen, T., Præbel, K., Hanebrekke, T., Ray, J. L., Fall, J., and Westgaard J. I. 2024. Predicting Trawl Catches Using Environmental DNA. ICES Journal of Marine Science: Journal Du Conseil, August, fsae097.

Hijmans, R. J. (2024). raster: Geographic Data Analysis and Modeling. https://rspatial.org/raster

Izquierdo-Serrano, M., Revuelta, O., Míguez-Lozano, R., Gozalbes, P., Ruiz-García, D., Raga, J. A., & Tomás, J. (2022). Assessment of the interactions between cetaceans and fisheries at the south of the Cetacean Migration Corridor and neighboring waters (Western Mediterranean). Frontiers in Marine Science, 9, 981638.

Jefferson, T. A., Weir, C. R., Anderson, R. C., Ballance, L. T., Kenney, R. D., & Kiszka, J. J. (2014). Global distribution of Risso’s dolphin Grampus griseus: a review and critical evaluation. Mammal Review, 44(1), 56–68.

Kaschner, K., Quick, N. J., Jewell, R., Williams, R., & Harris, C. M. (2012). Global coverage of cetacean line-transect surveys: status quo, data gaps and future challenges. PloS One, 7(9), e44075.

Kelly, R. P., Shelton, A. O., & Gallego, R. (2019). Understanding PCR Processes to Draw Meaningful Conclusions from Environmental DNA Studies. Scientific Reports, 9(1), 12133.

Kowarski, K. A., Delarue, J. J.-Y., Gaudet, B. J., & Martin, S. B. (2021). Automatic data selection for validation: A method to determine cetacean occurrence in large acoustic data sets. JASA Express Letters, 1(5), 051201.

Kuhn, M. (2008). Building Predictive Models inRUsing thecaretPackage. Journal of Statistical Software, 28(5), 1–26.

Lahoz-Monfort, J. J., Guillera-Arroita, G., & Tingley, R. (2016). Statistical approaches to account for false-positive errors in environmental DNA samples. Molecular Ecology Resources, 16(3), 673–685.

Lewin, W.-C., Weltersbach, M. S., Ferter, K., Hyder, K., Mugerza, E., Prellezo, R., Radford, Z., Zarauz, L., & Strehlow, H. V. (2019). Potential environmental impacts of recreational fishing on marine fish stocks and ecosystems. Reviews in Fisheries Science & Aquaculture, 27(3), 287–330.

Martin, M. (2011). Cutadapt removes adapter sequences from high-throughput sequencing reads. EMBnet.Journal, 17(1), 10–12.

Mathieu, C., Hermans, S. M., Lear, G., Buckley, T. R., Lee, K. C., & Buckley, H. L. (2020). A Systematic Review of Sources of Variability and Uncertainty in eDNA Data for Environmental Monitoring. Frontiers in Ecology and Evolution, 8. 10.3389/fevo.2020.00135

Merten, V., GEOMAR Helmholtz Centre for Ocean Research Kiel, Visser, F., & Hoving, H.-J. (2023). Deep-sea predator-prey dynamics revealed by biologging and eDNA analysis. Oceanography (Washington, D.C.). 10.5670/oceanog.2023.s1.31

Merten, V. J., Visser, F., & Hoving, H.-J. T. (2023). Deep-Sea Predator-Prey Dynamics Revealed by Biologging and eDNA Analysis. Oceanography, 36(1), 96–99.

Meynecke, J.-O., de Bie, J., Barraqueta, J.-L. M., Seyboth, E., Dey, S. P., Lee, S. B., Samanta, S., Vichi, M., Findlay, K., Roychoudhury, A., & Mackey, B. (2021). The role of environmental drivers in humpback whale distribution, movement and behavior: A review. Frontiers in Marine Science, 8, 720774.

Miller, D. L., Becker, E. A., Forney, K. A., Roberts, J. J., Cañadas, A., & Schick, R. S. (2022). Estimating uncertainty in density surface models. PeerJ, 10, e13950.

New England Biolabs. (n.d.). PCR Protocol for Phusion® High-Fidelity DNA Polymerase. Retrieved February 20, 2024, from https://www.neb.com/en-us/protocols/0001/01/01/pcr-protocol-m0530

Nisi, A. C., Welch, H., Brodie, S., Leiphardt, C., Rhodes, R., Hazen, E. L., Redfern, J. V., Branch, T. A., Barreto, A. S., Calambokidis, J., Clavelle, T., Dares, L., de Vos, A., Gero, S., Jackson, J. A., Kenney, R. D., Kroodsma, D., Leaper, R., McCauley, D. J., … Abrahms, B. (2024). Ship collision risk threatens whales across the world’s oceans. Science (New York, N.Y.), 386(6724), 870–875.

Pelayo-González, L., Herra-Miranda, D., Pacheco-Polanco, J. D., Guzmán, H. M., Goodman, S., & Oviedo, L. (2022). Decreases in encounter rate of endangered Northeast Pacific humpback whales in Southern Costa Rica: Possible changes in migration pattern due to warming events. Frontiers in Marine Science, 9. 10.3389/fmars.2022.927276

Phillips, E. M., Malick, M. J., Gauthier, S., Haltuch, M. A., Hunsicker, M. E., Parker-Stetter, S. L., & Thomas, R. E. (2023). The influence of temperature on Pacific hake co-occurrence with euphausiids in the California Current Ecosystem. Fisheries Oceanography, 32(3), 267–279.

Pikesley, S. K., Witt, M. J., Hardy, T., Loveridge, J., Loveridge, J., Williams, R., & Godley, B. J. (2012). Cetacean sightings and strandings: evidence for spatial and temporal trends? Journal of the Marine Biological Association of the United Kingdom. Marine Biological Association of the United Kingdom, 92(8), 1809–1820.

Pirotta, V., Grech, A., Jonsen, I. D., Laurance, W. F., & Harcourt, R. G. (2019). Consequences of global shipping traffic for marine giants. Frontiers in Ecology and the Environment, 17(1), 39–47.

Porter, T. M., & Hajibabaei, M. (2020). Putting COI Metabarcoding in Context: The Utility of Exact Sequence Variants (ESVs) in Biodiversity Analysis. Frontiers in Ecology and Evolution, 8. 10.3389/fevo.2020.00248

Poyntz-Wright, I. P., Harrison, X. A., Pedersen, S., & Tyler, C. R. (2024). Effectiveness of eDNA for monitoring riverine macroinvertebrates. The Science of the Total Environment, 941(173621), 173621.

Praca, E., & Gannier, A. (2008). Ecological niches of three teuthophageous odontocetes in the northwestern Mediterranean Sea. Ocean Science, 4(1), 49–59.

Qgis.org. (2024). QGIS Geographic Information System. Open Source Geospatial Foundation Project. https://qgis.org/

R Core Team. (2021). R: A Language and Environment for Statistical Computing. R Foundation for Statistical Computing. https://www.R-project.org/

Ramón-Laca, A., Wells, A., & Park, L. (2021). A workflow for the relative quantification of multiple fish species from oceanic water samples using environmental DNA (eDNA) to support large-scale fishery surveys. PloS One, 16(9), e0257773.

Rice, A., Debich, A. J., Širović, A., Oleson, E. M., Trickey, J. S., Varga, L. M., Wiggins, S. M., Hildebrand, J. A., & Baumann-Pickering, S. (2021). Cetacean occurrence offshore of Washington from long-term passive acoustic monitoring. Marine Biology, 168(8), 1–22.

Rockwood, R. C., Elliott, M. L., Saenz, B., Nur, N., & Jahncke, J. (2020). Modeling predator and prey hotspots: Management implications of baleen whale co-occurrence with krill in Central California. PloS One, 15(7), e0235603.

RStudio Team. (2020). RStudio: Integrated Development Environment for R. RStudio, PBC. http://www.rstudio.com/

Sahri, A., Herwata Putra, M. I., Kusuma Mustika, P. L., Kreb, D., & Murk, A. J. (2021). Cetacean habitat modelling to inform conservation management, marine spatial planning, and as a basis for anthropogenic threat mitigation in Indonesia. Ocean & Coastal Management, 205(105555), 105555.

Salvadeo, C. J., Lluch-Belda, D., Gómez-Gallardo, A., Urbán-Ramírez, J., & MacLeod, C. D. (2010). Climate change and a poleward shift in the distribution of the Pacific white-sided dolphin in the northeastern Pacific. Endangered Species Research, 11, 13–19.

Sambrook, J., Fritsch, E. F., & Maniatis, T. (1989). Molecular cloning: a laboratory manual (2d ed.). Cold Spring Harbor Laboratory.

Sbrocco, E. J., & Barber, P. H. (2013). MARSPEC: ocean climate layers for marine spatial ecology. Ecology, 94(4), 979–979.

Schorr, G., Falcone, E., Calambokidid, J., Hildebrand, J., & Oleson, E. (2007). Acoustic and visual monitoring for cetaceans along the outer Washington coast. https://calhoun.nps.edu/bitstream/handle/10945/704/NPS-OC-09-001.pdf?sequence=3

Shelton, A. O., Gold, Z. J., Jensen, A. J., D Agnese, E., Andruszkiewicz Allan, E., Van Cise, A., Gallego, R., Ramón-Laca, A., Garber-Yonts, M., Parsons, K., & Kelly, R. P. (2023). Toward quantitative metabarcoding. Ecology, 104(2), e3906.

Sing, T., Sander, O., Beerenwinkel, N., & Lengauer, T. (2005). ROCR: visualizing classifier performance in R. Bioinformatics (Oxford, England), 21(20), 3940–3941.

Stat, M., John, J., DiBattista, J. D., Newman, S. J., Bunce, M., & Harvey, E. S. (2018). Combined use of eDNA metabarcoding and video surveillance for the assessment of fish biodiversity. Conservation Biology: The Journal of the Society for Conservation Biology, 0(0), 1–10.

Szesciorka, A. R., Demer, D. A., Santora, J. A., Forney, K. A., & Moore, J. E. (2023). Multiscale relationships between humpback whales and forage species hotspots within a large marine ecosystem. Ecological Applications: A Publication of the Ecological Society of America, 33(2), e2794.

Thompson, S. A., Sydeman, W. J., Santora, J. A., Black, B. A., Suryan, R. M., Calambokidis, J., Peterson, W. T., & Bograd, S. J. (2012). Linking predators to seasonality of upwelling: Using food web indicators and path analysis to infer trophic connections. Progress in Oceanography, 101(1), 106–120.

Tixier, P., Lea, M.-A., Hindell, M. A., Welsford, D., Mazé, C., Gourguet, S., & Arnould, J. P. Y. (2021). When large marine predators feed on fisheries catches: Global patterns of the depredation conflict and directions for coexistence. Fish and Fisheries (Oxford, England), 22(1), 31–53.

Tobeña, M., Prieto, R., Machete, M., & Silva, M. A. (2016). Modeling the potential distribution and richness of cetaceans in the Azores from fisheries observer program data. Frontiers in Marine Science, 3, 202655.

United States. (1983). The Endangered Species Act as Amended by Public Law 97-304 (the Endangered Species Act Amendments of 1982). Washington :U.S. G.P.O. https://www.govinfo.gov/content/pkg/COMPS-3002/pdf/COMPS-3002.pdf

Visser, F., Merten, V. J., Bayer, T., Oudejans, M. G., de Jonge, D. S. W., Puebla, O., Reusch, T. B. H., Fuss, J., & Hoving, H. J. T. (2021). Deep-sea predator niche segregation revealed by combined cetacean biologging and eDNA analysis of cephalopod prey. Science Advances, 7(14). 10.1126/sciadv.abf5908

Vukelic, M., Mancini, F., Vukelic, D., & Carere, C. (2018). A cetacean monitoring system that integrates citizen science and satellite imagery. Rendiconti Lincei. Scienze Fisiche e Naturali, 29(1), 53–59.

Wang, Q., Garrity, G. M., Tiedje, J. M., & Cole, J. R. (2007). Naive Bayesian classifier for rapid assignment of rRNA sequences into the new bacterial taxonomy. Applied and Environmental Microbiology, 73(16), 5261–5267.

Wood, S. N. (2003). Thin plate regression splines. Journal of the Royal Statistical Society. Series B, Statistical Methodology, 65(1), 95–114.

Wood, S. N. (2008). Fast stable direct fitting and smoothness selection for generalized additive models. Journal of the Royal Statistical Society. Series B, Statistical Methodology, 70(3), 495–518.

Wood, S. N. (2017). Generalized additive models: An introduction with R, second edition. CRC Press.

Wood, S. N. (2023). mgcv: Mixed GAM Computation Vehicle with Automatic Smoothness Estimation (Version 1.9-1). https://cran.r-project.org/web/packages/mgcv/index.html

Wright, D. L., Witteveen, B., Wynne, K., & Horstmann-Dehn, L. (2015). Evidence of two subaggregations of humpback whales on the Kodiak, Alaska, feeding ground revealed from stable isotope analysis. Marine Mammal Science, 31(4), 1378–1400.

