## Supplementary Material for "Modeling cetacean eDNA distribution along the Washington coast using metabarcoding from opportunistic samples and generalized additive models"

**Table S1. Primers and adapters used for library construction using eDNA samples and targeting the D-loop mitochondrial region of marine mammals (Baker et al., 2018).**

| Primer | Sequence |
|---|---|
| Dlp1.5-H | 5′-TCACCCAAAGCTGRARTTCTA-3′ |
| Oordlp4 | 5′-GCGGGTTGCTGGTTTCACG-3′ |
| | P5 overhang: 5′-TCGTCGGCAGCGTCAGATGTGTATAAGAGACAG-forward-primer-sequence-3′ |
| | P7 overhang: 5′-GTCTCGTGGGCTCGGAGATGTGTATAAGAGACAG-reverse-primer-sequence-3′ |

**Table S2. Environmental covariates included in GAM models.**

| Dataset | Code | Description | Reference | Resolution | units | year | layer_url |
|---|---|---|---|---|---|---|---|
| MARSPEC | bathy | Bathymetry | https://esapubs.org/archive/ecol/E094/086/metadata.php | 0.083 | meters | 2009 | https://www.lifewatch.be/sdmpredictors/MS_bathy_5m_lonlat.tif |
| MARSPEC | dist_shore | Distance to shore | https://esapubs.org/archive/ecol/E094/086/metadata.php | 0.083 | kilometers | 2009 | https://www.lifewatch.be/sdmpredictors/MS_biogeo05_dist_shore_5m_lonlat.tif |
| MARSPEC | slope | Bathymetric slope | https://esapubs.org/archive/ecol/E094/086/metadata.php | 0.083 | degrees | 2009 | https://www.lifewatch.be/sdmpredictors/MS_biogeo06_bathy_slope_5m_lonlat.tif |
| COPERNICUS | SST | Sea water potential temperature | https://catalogue.marine.copernicus.eu/documents/PUM/CMEMS-GLO-PUM-001-030.pdf | 0.083 | °C | 2019 | https://data.marine.copernicus.eu/product/GLOBAL_MULTIYEAR_PHY_001_030/description |

**Table S3. Summary of the number of sequencing reads per technical replicate at each step of the bioinformatic processing pipeline.**

| | input | filtered | denoisedF | denoisedR | merged | nonchim | length_filter |
|---|---|---|---|---|---|---|---|
| **replicate1** | 9,465,852 | 4,891,072 | 4,881,252 | 4,883,762 | 3,046,614 | 2,878,836 | 1,308,747 |
| **replicate2** | 9,857,378 | 4,662,530 | 4,654,513 | 4,655,286 | 3,017,586 | 2,773,095 | 1,386,534 |
| **replicate3** | 8,534,845 | 3,976,171 | 3,966,009 | 3,968,152 | 2,601,096 | 2,419,585 | 1,138,765 |
| average | 9,286,025 | 4,509,924 | 4,500,591 | 4,502,400 | 2,888,432 | 2,690,505 | 1,278,015 |

**Table S4. GAMs with an AIC difference <2 for the three most frequently detected cetacean species along the coast of Washington, incorporating environmental variables such as bathymetry (bathy), distance to shore (dist_shore), latitude (lat), longitude (lon), slope, and seawater temperature (SST), along with statistical output parameters including AIC, ICOMP, df, logLink, deltaAIC, and weight. Models are ordered by AIC value.**

| Pacific white-sided dolphin | | | | | | | | | | | | |
|---|---|---|---|---|---|---|---|---|---|---|---|---|
| (Int) | s(bth,"ts") | s(dist_shore,"ts") | s(lat,"ts") | s(lon,"ts") | s(slope,"ts") | s(SST,"ts") | AIC | ICOMP | df | logLik | delta | weight |
| -2.9 | | + | | + | | | 29.1 | 77.94 | 3 | -11.52 | 0 | 0.09 |
| -2.9 | | + | + | + | | | 29.1 | 216.03 | 3 | -11.52 | 0 | 0.09 |
| -2.9 | | + | | + | | + | 29.1 | 195.61 | 3 | -11.52 | 0 | 0.09 |
| -2.9 | | + | | + | + | | 29.1 | 215.05 | 3 | -11.52 | 0 | 0.09 |
| -2.72 | | + | | | | + | 29.48 | 74.79 | 3 | -11.63 | 0.38 | 0.07 |
| -2.72 | | + | + | | | + | 29.48 | 207.63 | 3 | -11.63 | 0.38 | 0.07 |
| -2.72 | | + | | | + | + | 29.48 | 215.29 | 3 | -11.63 | 0.38 | 0.07 |
| -2.82 | | + | + | | | | 30.13 | 88.68 | 3 | -12.06 | 1.03 | 0.05 |
| -2.82 | | + | + | | + | | 30.14 | 224.52 | 3 | -12.06 | 1.03 | 0.05 |
| -2.26 | + | | | + | | | 30.98 | 71.35 | 3 | -12.39 | 1.88 | 0.03 |
| -2.26 | + | | + | + | | | 30.98 | 196.5 | 3 | -12.39 | 1.88 | 0.03 |
| -2.26 | + | | | + | + | | 30.98 | 199.54 | 3 | -12.39 | 1.88 | 0.03 |
| -2.95 | | | | + | | + | 30.98 | 70.88 | 3 | -12.34 | 1.88 | 0.03 |
| -2.95 | | | + | + | | + | 30.98 | 196.73 | 3 | -12.34 | 1.88 | 0.03 |
| -2.95 | | | | + | + | + | 30.98 | 209.45 | 3 | -12.34 | 1.88 | 0.03 |
| -2.64 | | + | | | + | | 31.11 | 186.15 | 2 | -13.48 | 2.01 | 0.03 |

| | | | | | | | | | | | | |
|---|---|---|---|---|---|---|---|---|---|---|---|---|
| -2.49 | + | | | + | | + | 31.31 | 101.2 | 3 | -11.93 | 2.2 | 0.03 |
| -2.3 | | | + | + | | | 31.7 | 74.7 | 3 | -12.48 | 2.6 | 0.02 |
| -2.3 | | | + | + | + | | 31.7 | 213.52 | 3 | -12.48 | 2.6 | 0.02 |
| -1.93 | + | | + | | | | 35.49 | 83.06 | 3 | -14.63 | 6.39 | 0 |
| -1.93 | + | | + | | + | | 35.49 | 208.36 | 3 | -14.63 | 6.39 | 0 |
| -1.93 | + | | + | | | + | 35.49 | 190.94 | 3 | -14.63 | 6.39 | 0 |
| -1.6 | | | | + | + | | 37.02 | 184.6 | 2 | -16.44 | 7.92 | 0 |
| -1.78 | + | | | | | + | 37.58 | 180.79 | 2 | -16.69 | 8.47 | 0 |
| -1.78 | + | | | | + | + | 37.58 | 315.39 | 2 | -16.69 | 8.47 | 0 |
| -1.78 | + | | | | + | | 37.58 | 171.91 | 2 | -16.69 | 8.47 | 0 |
| -2.11 | | | + | | | + | 39.49 | 83.83 | 3 | -16.51 | 10.38 | 0 |
| -2.11 | | | + | | + | + | 39.49 | 224.98 | 3 | -16.51 | 10.38 | 0 |
| -1.25 | | | | | + | + | 47.99 | 195.1 | 1 | -22.08 | 18.88 | 0 |
| -1.19 | | | + | | + | | 48.88 | 206.14 | 1 | -23.15 | 19.78 | 0 |

**Humback whale**

| (Int) | s(bth,"ts") | s(dist_shore,"ts") | s(lat,"ts") | s(lon,"ts") | s(slope,"ts") | s(SST,"ts") | AIC | ICOMP | df | logLik | delta | weight |
|---|---|---|---|---|---|---|---|---|---|---|---|---|
| -1.97 | + | | | | | + | 35.45 | 74.24 | 3 | -14.64 | 0 | 0.19 |
| -1.97 | + | | | + | | + | 35.45 | 209.59 | 3 | -14.64 | 0 | 0.19 |
| -1.97 | + | | + | | | + | 35.45 | 207.2 | 3 | -14.64 | 0 | 0.19 |
| -1.93 | + | | | | + | + | 35.87 | 108.74 | 3 | -14.31 | 0.42 | 0.15 |

| | | | | | | | | | | | | |
|---|---|---|---|---|---|---|---|---|---|---|---|---|
| -1.76 | | | | | + | + | 39.94 | 84.29 | 2 | -17.07 | 4.48 | 0.02 |
| -1.76 | | | + | | + | + | 39.94 | 213.89 | 2 | -17.07 | 4.48 | 0.02 |
| -1.76 | | + | | | + | + | 39.94 | 194.34 | 2 | -17.07 | 4.48 | 0.02 |
| -1.76 | | | | + | + | + | 39.94 | 196.18 | 2 | -17.07 | 4.48 | 0.02 |
| -1.68 | + | | | + | | | 40.22 | 186.92 | 1 | -18.12 | 4.77 | 0.02 |
| -1.68 | + | | + | + | | | 40.22 | 296.2 | 1 | -18.12 | 4.77 | 0.02 |
| -1.68 | + | | + | | | | 40.22 | 171.65 | 1 | -18.12 | 4.77 | 0.02 |
| -1.68 | | + | | | + | | 40.6 | 190.08 | 1 | -18.31 | 5.15 | 0.01 |
| -1.68 | | + | + | | + | | 40.6 | 328.5 | 1 | -18.31 | 5.15 | 0.01 |
| -1.68 | | | | + | + | | 40.6 | 178.29 | 1 | -18.31 | 5.15 | 0.01 |
| -1.68 | | | + | + | + | | 40.6 | 300.77 | 1 | -18.31 | 5.15 | 0.01 |
| -1.68 | | + | | + | + | | 40.6 | 303.48 | 1 | -18.31 | 5.15 | 0.01 |
| -1.68 | | | + | | + | | 40.6 | 170.37 | 1 | -18.31 | 5.15 | 0.01 |
| -1.68 | + | | | | + | | 40.68 | 89.63 | 2 | -17.68 | 5.23 | 0.01 |
| -1.68 | + | | | + | + | | 40.68 | 208.43 | 2 | -17.68 | 5.23 | 0.01 |
| -1.68 | + | | + | | + | | 40.69 | 167.6 | 2 | -17.67 | 5.24 | 0.01 |
| -1.48 | | | + | | | + | 43.32 | 301.49 | 1 | -20.66 | 7.87 | 0 |
| -1.48 | | + | + | | | | 43.37 | 220.26 | 1 | -20.63 | 7.92 | 0 |
| -1.5 | | + | + | + | | | 43.51 | 330.38 | 1 | -20.02 | 8.06 | 0 |
| -1.5 | | + | | + | | + | 43.51 | 344.74 | 1 | -20.02 | 8.06 | 0 |
| -1.5 | | | + | + | | + | 43.51 | 336.02 | 1 | -20.02 | 8.06 | 0 |

| | | | | | | | | | | | | |
|---|---|---|---|---|---|---|---|---|---|---|---|---|
| -1.5 | | + | | + | | | 43.51 | 189.5 | 1 | -20.02 | 8.06 | 0 |
| -1.5 | | | + | + | | | 43.51 | 192.67 | 1 | -20.02 | 8.06 | 0 |
| -1.5 | | | | + | | + | 43.51 | 186.62 | 1 | -20.02 | 8.06 | 0 |
| -1.51 | | + | | | | + | 43.62 | 95.85 | 2 | -19.21 | 8.17 | 0 |
| -1.51 | | + | + | | | + | 43.62 | 223.32 | 2 | -19.21 | 8.17 | 0 |

**Risso's dolphin**

| (Int) | s(bth,"ts") | s(dist_shore,"ts") | s(lat,"ts") | s(lon,"ts") | s(slope,"ts") | s(SST,"ts") | AIC | ICOMP | df | logLik | delta | weight |
|---|---|---|---|---|---|---|---|---|---|---|---|---|
| -2.26 | | | | + | + | | 32.73 | 76.18 | 2 | -13.4 | 0 | 0.06 |
| -2.26 | | | + | + | + | | 32.73 | 204.62 | 2 | -13.4 | 0 | 0.06 |
| -2.26 | | | | + | + | + | 32.73 | 184.68 | 2 | -13.4 | 0 | 0.06 |
| -2.26 | + | | | + | + | | 32.73 | 180.63 | 2 | -13.4 | 0 | 0.06 |
| -2.35 | | + | | | + | | 33.08 | 80.52 | 2 | -13.65 | 0.35 | 0.05 |
| -2.35 | | + | | | + | + | 33.08 | 192.65 | 2 | -13.65 | 0.35 | 0.05 |
| -2.42 | | + | | + | + | | 33.1 | 108.73 | 3 | -12.89 | 0.37 | 0.05 |
| -2.38 | | + | + | | + | | 33.3 | 122.02 | 3 | -13.26 | 0.57 | 0.05 |
| -2.2 | | + | + | | | + | 33.41 | 315.22 | 2 | -14.69 | 0.68 | 0.05 |
| -2.2 | | + | | | | + | 33.41 | 171.96 | 2 | -14.69 | 0.68 | 0.05 |
| -2.2 | | + | + | | | | 33.41 | 182.46 | 2 | -14.69 | 0.68 | 0.05 |
| -2.2 | | + | | + | | | 33.84 | 92.42 | 2 | -14.38 | 1.11 | 0.04 |
| -2.2 | | + | | + | | + | 33.84 | 213.68 | 2 | -14.38 | 1.11 | 0.04 |

| | | | | | | | | | | | | |
|---|---|---|---|---|---|---|---|---|---|---|---|---|
| -2.2 | | + | + | + | | | 33.84 | 222.62 | 2 | -14.38 | 1.11 | 0.04 |
| -1.98 | | | + | + | | | 34.71 | 183.44 | 1 | -15.38 | 1.98 | 0.02 |
| -2.11 | | | | + | | + | 34.78 | 83.08 | 2 | -14.6 | 2.05 | 0.02 |
| -2.11 | | | + | + | | + | 34.78 | 204.52 | 2 | -14.6 | 2.05 | 0.02 |
| -2.11 | + | | | + | | + | 34.78 | 193.37 | 2 | -14.6 | 2.05 | 0.02 |
| -2.02 | + | | | + | | | 35.12 | 89.14 | 2 | -14.91 | 2.4 | 0.02 |
| -2.02 | + | | + | + | | | 35.12 | 209.76 | 2 | -14.91 | 2.4 | 0.02 |
| -1.97 | | | | | + | + | 35.14 | 178.69 | 1 | -15.6 | 2.41 | 0.02 |
| -1.98 | | | + | | + | | 35.33 | 99.59 | 2 | -15.37 | 2.61 | 0.02 |
| -1.98 | | | + | | + | + | 35.33 | 211.98 | 2 | -15.37 | 2.61 | 0.02 |
| -2.04 | + | | | | + | | 35.39 | 85.25 | 2 | -14.98 | 2.66 | 0.02 |
| -2.04 | + | | | | + | + | 35.39 | 208.47 | 2 | -14.98 | 2.66 | 0.02 |
| -2.09 | + | | + | | + | | 35.52 | 118.88 | 3 | -14.37 | 2.79 | 0.02 |
| -1.94 | + | | | | | + | 35.67 | 180.26 | 1 | -15.9 | 2.94 | 0.01 |
| -1.94 | + | | + | | | + | 35.67 | 299.74 | 1 | -15.9 | 2.94 | 0.01 |
| -1.94 | + | | + | | | | 35.67 | 177.77 | 1 | -15.9 | 2.94 | 0.01 |
| -1.94 | | | + | | | + | 36.94 | 90.46 | 2 | -15.76 | 4.21 | 0.01 |

**Figure S5. Partial effects plot showing the relationship between a) longitude and distance to shore and the occurrence of eDNA for Pacific white-sided dolphin, b) bathymetry and sea water temperature and the occurrence of eDNA for humpback whale, and c) longitude and slope and the occurrence of eDNA for Risso's dolphin. The plots illustrate the predicted probability of eDNA occurrence as a function of each covariate while holding all other predictors constant at their mean values.**

**a)**

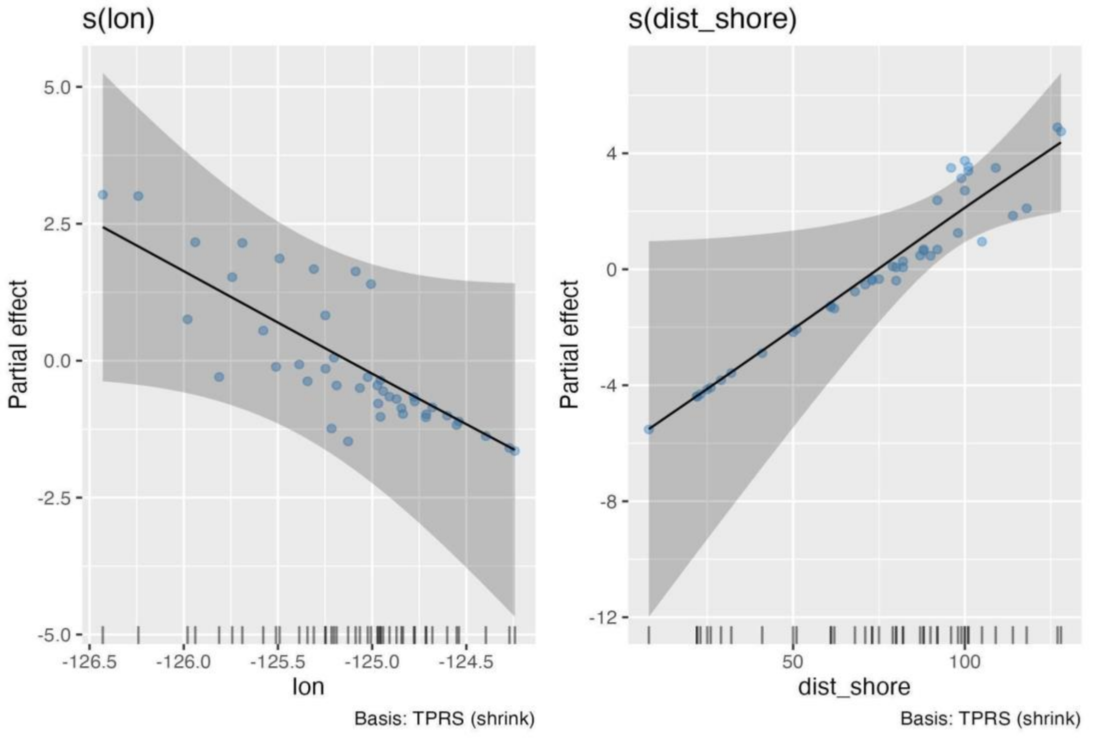

**b)**

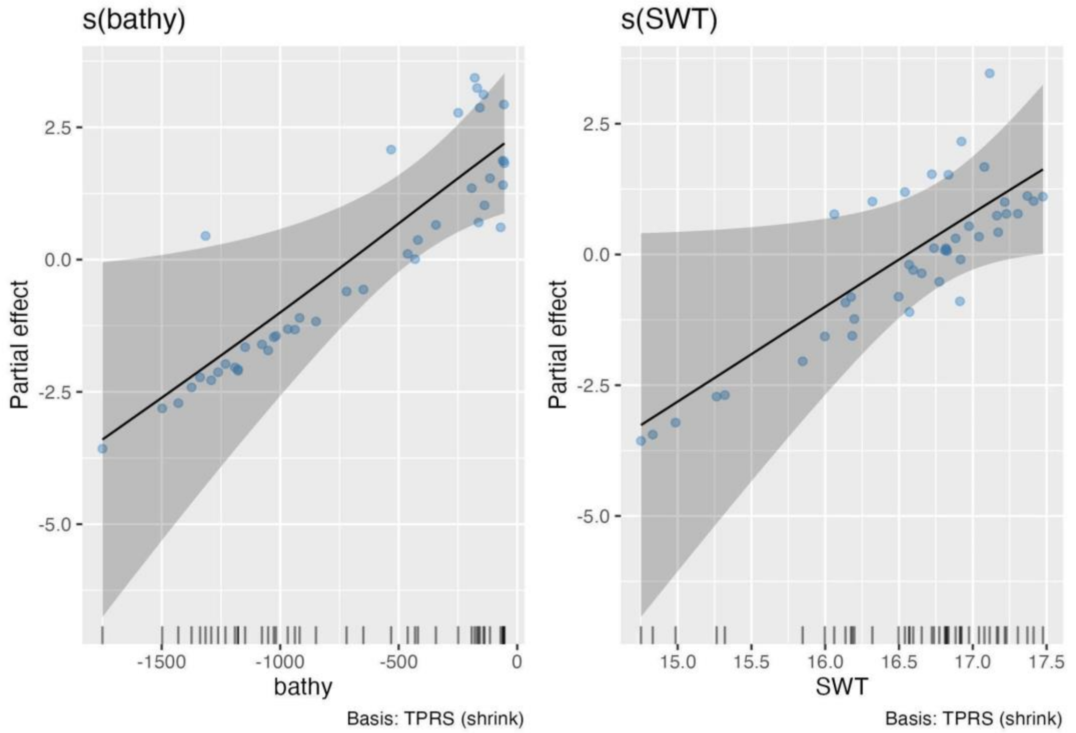
s(bathy)
Partial effect
2.5
0.0
-2.5
-5.0
-1500
-1000
-500
0
bathy
Basis: TPRS (shrink)
s(SWT)
Partial effect
2.5
0.0
-2.5
-5.0
15.0
15.5
16.0
16.5
17.0
17.5
SWT
Basis: TPRS (shrink)

**c)**

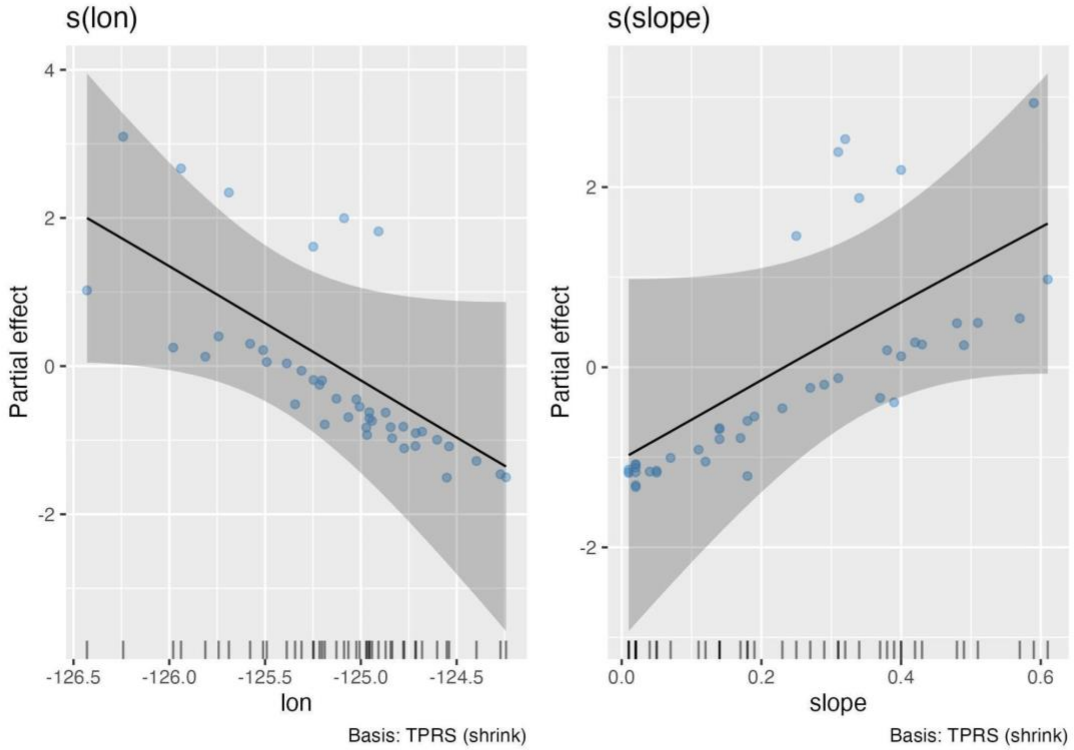
s(lon)
s(slope)
Partial effect
Partial effect
4
2
0
-2
-2
-126.5
-126.0
-125.5
-125.0
-124.5
0.0
0.2
0.4
0.6
lon
slope
Basis: TPRS (shrink)
Basis: TPRS (shrink)
